# The Ca^2+^ Permeation Mechanism of the Ryanodine Receptor Revealed by a Multi-Site Ion Model

**DOI:** 10.1101/682328

**Authors:** Aihua Zhang, Hua Yu, Chunhong Liu, Chen Song

## Abstract

The ryanodine receptors (RyR) are ion channels responsible for the release of Ca^2+^ from the sarco/endoplasmic reticulum and play a crucial role in the precise control of Ca^2+^ concentration in the cytosol. The detailed permeation mechanism of Ca^2+^ through RyR is still elusive. By using molecular dynamics simulations with a specially designed Ca^2+^ model, here we show that multiple Ca^2+^ accumulate in the upper selectivity filter of RyR1, but only one Ca^2+^ can enter and translocate in the narrow pore at a time. The Ca^2+^ is nearly fully hydrated during the whole permeation process, with the first solvation shell intact even at the narrowest constrict sites of the selectivity filter and gate. These results present a one-at-a-time permeation pattern for the hydrated ions, which is distinct from the fully/partially dehydrated knock-on permeation in K^+^ and Na^+^ channels and uncovers the underlying reason for the high permeability and low selectivity of the RyR channels.

## Introduction

As an essential messenger in cells, calcium ions (Ca^2+^) regulate many physiological processes, including neurotransmitter release, muscle contraction, and hormones secretion^1^. The concentration of Ca^2+^ in the cytoplasm and organelles is precisely controlled by multiple calcium channels, including the voltage-gated calcium channels in cell membranes and the ryanodine receptors (RyR) in the endoplasmic reticulum (ER) membrane. Ca^2+^ also induces conformational changes of a wide range of Ca^2+^-interacting proteins, such as calmodulin and Ca^2+^-activated ion channels, to trigger downstream signal transduction^2^. Although the pathways of calcium signaling are extensively studied, the molecular interaction details between calcium and proteins have yet to be fully elucidated.

To study the detailed interactions between ions and proteins, we can often use molecular dynamics (MD) simulations to provide microscopic and quantitative insights, thereby obtaining the specific functional mechanisms of the relevant proteins^3–6^. However, the conventional models of Ca^2+^ are far from accurate in calculating the interaction energies between Ca^2+^ and proteins^7–9^, therefore inadequate to study the precise Ca^2+^-protein interactions. As a consequence, K^+^ and Na^+^ channels have been widely studied by MD simulations, and their detailed permeation mechanisms were revealed^10–17^, but computational studies of Ca^2+^ channels are rather limited. Notably, several structures of Ca^2+^ channels were resolved recently^18–21^, which provided a solid basis to study their detailed function mechanism further. In particular, the open-state RyR1 channel provides an excellent opportunity for studying Ca^2+^ permeation and selectivity^21^, which makes a reliable Ca^2+^ model even more desirable.

There have been enormous efforts in trying to develop a more accurate Ca^2+^ model. The polarizable force field is theoretically appealing^9^, but its implementation and validation still need further work before being widely accepted and utilized for membrane protein simulations. Kohagen et al. proposed to scale the partial charges on the Ca^2+^ to account for the charge transfer and polarization^22,23^. Another strategy is to represent an ion by distributing electrostatic and Lennard-Jones (LJ) interactions on multiple sites, which can introduce a much larger parameter space and therefore make the ion model more tailorable^24–26^. Unfortunately, none of the existing Ca^2+^ models in the non-polarizable classical force field are able to describe the interactions between Ca^2+^ and protein quantitatively. Therefore, in the present work, we developed a new multi-site Ca^2+^ model particularly optimized for Ca^2+^-protein interaction (inset of Fig. 1), and then utilize this model to study the detailed Ca^2+^ permeation mechanism through the RyR1 channel, which showed distinct features from the widely studied K^+^ and Na^+^ channels.

**Figure 1.**
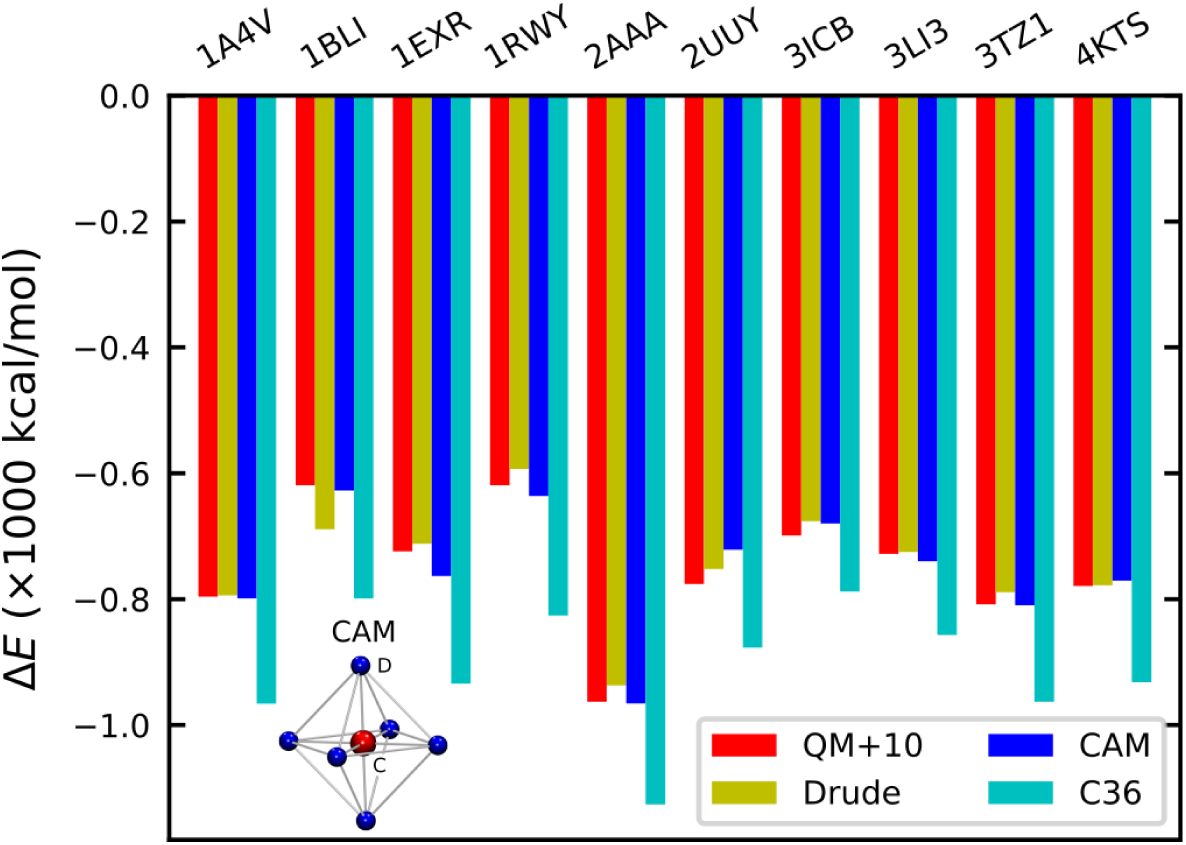
The calcium-protein binding energies calculated with different methods. Our multi-site Ca^2+^ model (CAM) consists of a central atom (C) and six dummy atoms (D) located at the vertices of an octahedron, as shown with the inset.

## Results

### The multi-site Ca^2+^ model behaves well in both water and protein

We designed a seven-site ion model, as shown in the inset of Fig. 1. There are six adjustable parameters, including *b*_*CD*_, *Q*_*C*_, *ε*_*C*_, *σ*_*C*_, *ε*_*D*_, and *σ*_*D*_, where *b*_*CD*_ is the distance between dummy atoms and the central atom, *Q*_*C*_ is the charge on the central atom, and the *ε*’s and *σ*’s are the LJ parameters of the central (C) and dummy (D) atoms, respectively. We further distinguish the LJ interactions of Ca^2+^ with water and non-water by replacing (*ε*_*C*_, *σ*_*C*_) with two sets of 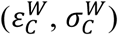 and 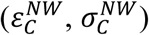, respectively. The charges on all the dummy atoms are the same and determined so that the total charge of the model is +2 *e*. By adjusting the aforementioned parameters, we obtained a Ca^2+^ model that can quantitatively reproduce the energetical and dynamic properties of Ca^2+^ in water as well as the Ca^2+^-protein interactions. The resulting Ca^2+^ properties in water are shown in Table 1. As can be seen, the hydration free energy (Δ*G*_*h*_), the first-peak position of the radial distribution function of water around Ca^2+^ (*R*_1_), and the number of coordinated water molecules in the first solvent shell (*N*_*C*_) have all reached the target values of experiments. In addition, the residence time of water molecules in the first solvation shell (*τ*_*R*_) can be optimized to below 100 ps, which solved the common problem of Ca^2+^ being too sticky. As there is no solid experimental data about the exact residence time of water, there may be still some room for further improvement. Nonetheless, our model shows that the water residence time in the first solvation shell can be optimized to a considerable extent with the multi-site model, and short residence time is consistent with a previous systematic study^27^.

**Table 1.** The target properties of Ca^2+^ in water and the computed property values from the simulations with our Ca^2+^ model.

| | $\Delta G_h$ (kJ/mol) | $R_1$ (nm) | $N_C$ | $\tau_R$ (ps) |
| --- | --- | --- | --- | --- |
| Target | -1504.0 | 0.242 | 7.0 | ~100 |
| Computed | -1503.9 | 0.242 | 7.0 | 75 |

With our model, the calculated binding energies of Ca^2+^ and proteins were also improved to a large extent (Fig. 1). The default Ca^2+^ parameters of the CHARMM force field (C36) led to a significant overestimation of ∼150–200 kcal/mol^9^, while the average binding-energy discrepancies for ten selected proteins were 6.6 kcal/mol for the Drude polarizable model and -0.2 kcal/mol for our multi-site model, respectively. Therefore, our model is comparable to the quantum mechanics (QM) and polarizable Drude model in calculating the Ca^2+^-protein binding energies, which is of crucial importance in simulating Ca^2+^-protein interactions and Ca^2+^ permeation through ion channels.

### The permeability of the open-state RyR1

The conventional ion models generally work well in studying ion-protein interactions for K^+^ or Na^+^, but fail consistently for Ca^2+^ ions^9^, and therefore no calcium permeation was observed in previous MD studies of Ca^2+^ channels^28^. We performed MD simulations on the type-1 ryanodine receptor (RyR1), an intracellular calcium release channel required for skeletal muscle contraction, with our Ca^2+^ model. The open-state structure of RyR1 (PDB ID: 5TAL) was obtained from des Georges et al.’s work^21^. The simulation systems consist of the pore domain of RyR1 embedded in a lipid bilayer of 1-Palmitoyl-2-oleoyl-sn-glycerol-3-phosphocholine (POPC) and an aqueous solution of either 150 mM Ca^2+^ or 250 mM K^+^ (Fig. 2a). The protein was restrained to the open-state crystal structure, and a transmembrane potential of 100 mV was applied along the direction from the sarcoplasmic reticulum (SR) lumen to the cytosol.

**Figure 2.**
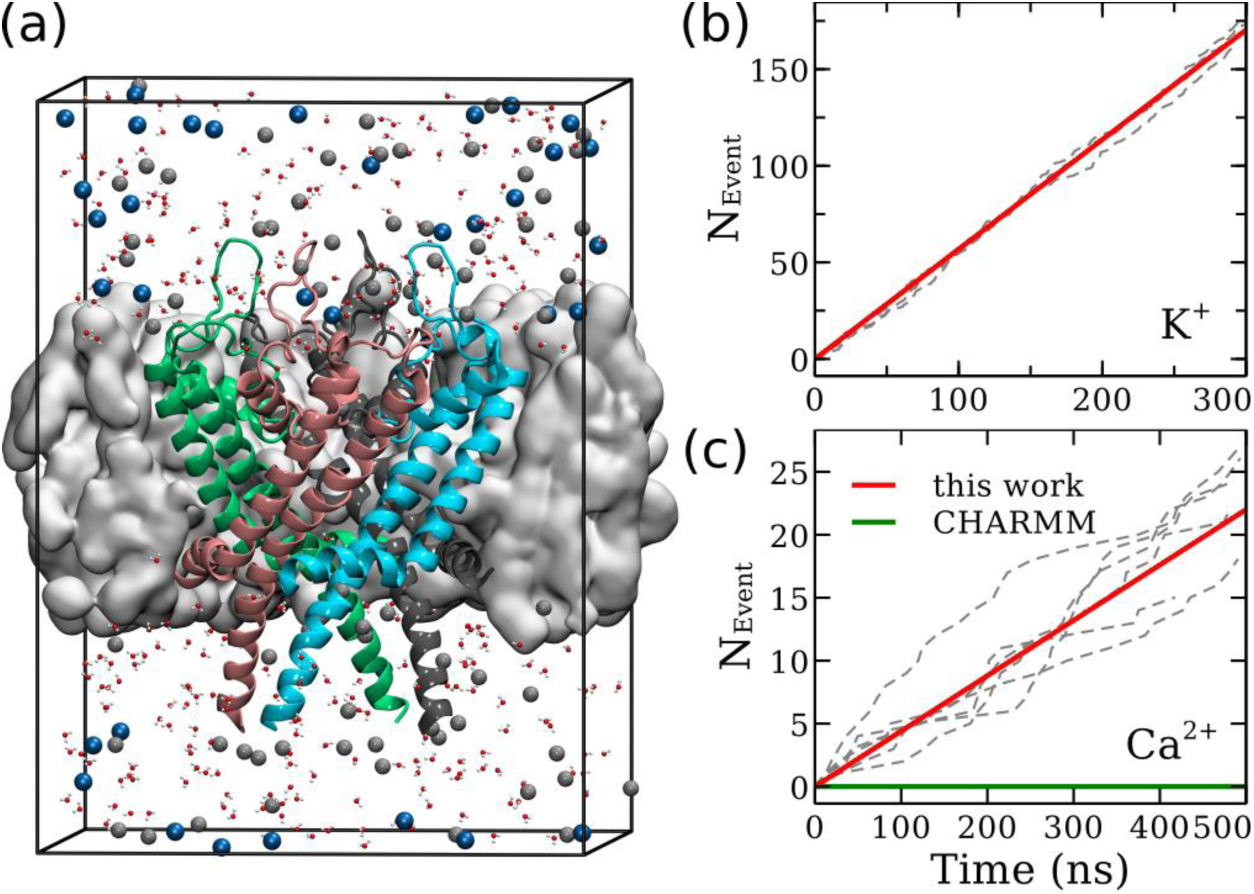
(a) The simulation system of Ca^2+^ (gray spheres) permeation through the RyR1 channel. The pore domain of RyR1 (cartoon) is embedded in a POPC membrane (gray surface). (b) The cumulative number of K^+^ ions permeating through the RyR1 channel as a function of simulation time. The dashed gray lines correspond to three independent simulation trajectories, and the solid red line corresponds to the average conductance. The conductance calculated from these trajectories is 910 ± 39 pS. (c) Same as (b) but for Ca^2+^ ions. The conductance from six trajectories is 141 ± 30 pS with our Ca^2+^ model.

We first studied the permeation of K^+^ through RyR1 as a validation. Three independent 300-ns MD simulations were conducted and generated enough permeation events (∼500) for statistical analysis (Fig. 2b). The conductance was calculated to be 910 ± 39 pS, which was in good agreement with the experimentally measured conductance of ∼850 pS with the same ion concentration and indicates that the RyR1 structure under study is indeed in its open state and that the K^+^ model in the CHARMM force field is reasonably accurate in describing the interactions between ions and proteins. However, our MD simulations of Ca^2+^ permeation through the same open-state RyR1 showed that the channel is not permeable to Ca^2+^ at all with the default Ca^2+^ parameter of CHARMM. Not a single permeation event was observed in three 500-ns MD trajectories (Fig. 2c). Since the channel is in its open state, this indicated that the default Ca^2+^ model gave us qualitatively wrong simulation results here, as also observed by another recent study^28^. A close inspection of the MD trajectories showed that the Ca^2+^ ions were tightly bound to the protein, again confirming that the binding affinity between Ca^2+^ and protein is too strong with the conventional Ca^2+^ model. In contrast, when our Ca^2+^ model was used for the same simulations, we observed continuous Ca^2+^ permeation as expected (Fig. 2c). The conductance calculated from six 500-ns trajectories is 141 ± 30 pS, which agrees reasonably well with the experimental value of ∼172 pS with the same ion concentration^29^. Therefore, we believe that our multi-site Ca^2+^ model is more accurate in studying the permeation behavior of the Ca^2+^ channel.

### The Ca^2+^ binding sites in the pore region

We performed an 800-ns simulation without the transmembrane potential to identify the Ca^2+^ binding sites in RyR1. The contour plot of the Ca^2+^ density, ρ(R, z), is presented in Fig. 3a, from which two major binding sites in the luminal vestibule (L) and above the selectivity filter constriction (S), and one minor binding site at the gate constriction (G) can be identified within the transmembrane pore. The selectivity filter and gate constrictions are located near the residues G4894 and Q4933, respectively (Fig. 3b). The corresponding positions in the contour plot are indicated by solid lines labeled with SF and GT in Fig. 3a. The isosurfaces of probability density corresponding to these binding sites are shown in Fig. 3b. By calculating the residence time of carboxylate oxygen of negatively charged residues within a sphere with a radius of 5 Å around the binding site L, we identified that the binding site L is formed by the interaction of Ca^2+^ ions with D4899, E4900, and D4903 residues (Fig. 3b), which agrees well with previous experimental and computational studies^28,30^. The large probability of finding Ca^2+^ in the selectivity filter also indicates that the Ca^2+^ can easily accumulate around the luminal vestibule and move into the upper filter, and the rate-determining step of permeation is the process of passing through the lower selectivity filter or gate constrictions. In addition, a continuous cytosolic binding region was found near residues D4938, E4942, and D4945, which interact with the permeating Ca^2+^ and may influence the ion permeability as well. This is consistent with a previous experimental study showing that D4938 and D4945 determines the ion flux and selectivity^31^.

**Figure 3.**
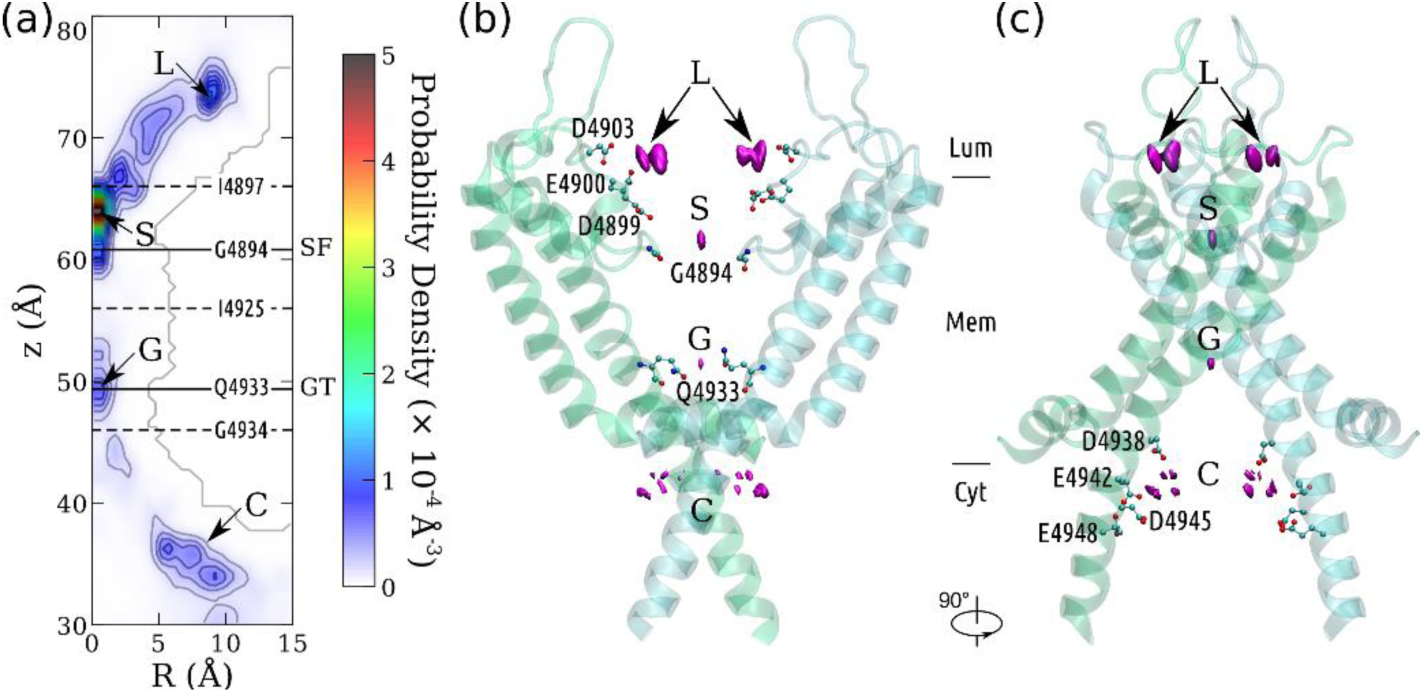
The Ca^2+^ binding sites in the RyR1 channel. (a) The contour plot of Ca^2+^ density on the R-z plane around RyR1. Four binding sites within the pore are designated as L, S, G, and C. The positions of the selectivity filter (SF) and gate (GT) constrictions are indicated with solid lines. The ion channel is divided into chambers by the dashed lines at the density saddles. The pore residues in close proximity are labeled on these lines. (b) & (c) Side views of the RyR1 channel. Only two chains are shown for clarity. The bottleneck residues (GLY-4894 at SF and GLN-4933 at GT), and the negatively charged residues at the binding sites L and C are shown as ball-and-sticks.

### The Ca^2+^ ions are fully hydrated during permeation

We calculated the number of oxygen atoms coordinated with the permeating Ca^2+^ and monitored from which residues these oxygen atoms were (water or protein). As shown in Fig. 4a, the open-state RyR pore is relatively wide compared to K^+^ and Na^+^ channels. As the first solvation shell is around 2.4 Å from the Ca^2+^ and the radius of a water molecule is usually considered to be 1.4 Å, we consider the pore region with a radius less than 4.0 Å to be the narrow pore region that contains the rate-limiting constriction sites. Interestingly, the average number of oxygen atoms coordinated with the Ca^2+^ was almost constant during the permeation process of Ca^2+^, as shown in Fig. 4a (right panel), and nearly all of these oxygen atoms were from water molecules within the narrow pore region. This clearly indicates that the permeating Ca^2+^ ions do not need to dehydrate when permeating through the open-state pore, and therefore the first solvation shell was intact in the narrow pore region. On the other hand, in the wide upper selectivity filter, one of the water molecules coordinated with Ca^2+^ can be replaced by the oxygen from E4900 (Fig. 4a), simply because the strong electrostatic attraction between Ca^2+^ and E4900 pushed one of the coordinated water molecules away. A similar phenomenon was observed near D4938. It should be noted that, the water replacement at these sites are not caused by the steric dehydration when ions passing through a narrow pore as observed in K^+^ and Na^+^ channels, but rather due to strong electrostatic attraction from negatively charged residues in a wide vestibule (r > 5 Å), and therefore should not be considered as dehydration due to the permeation.

**Figure 4.**
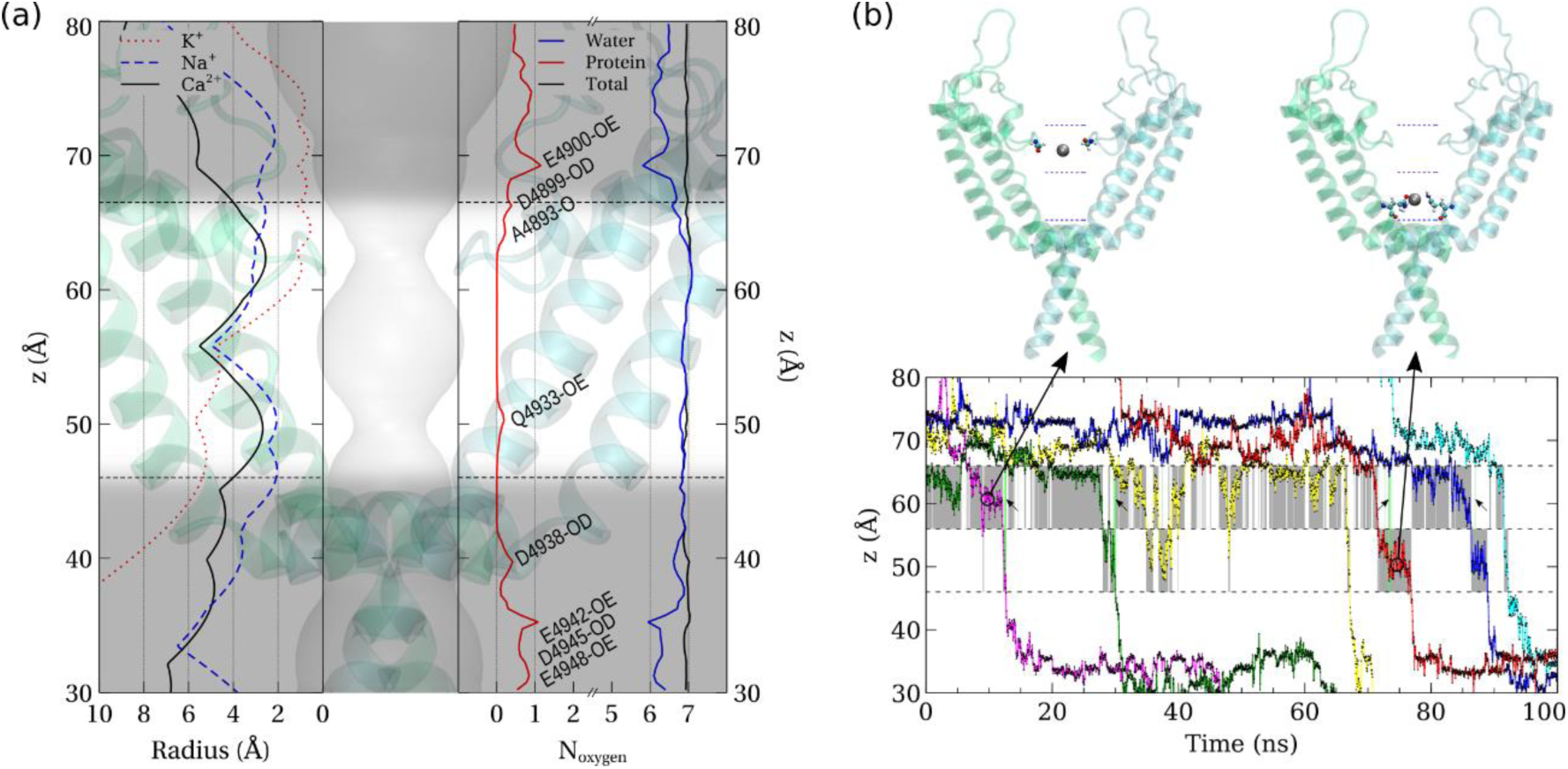
(a) The left panel shows the pore radii along the pore axis of the open-state RyR1 (black solid), K^+^ channel (red dotted), and Na^+^ channel (blue dashed). The right panel shows the total number (black) of coordinated oxygen atoms around the calcium ions within the pore, and the contributions from protein (red) and water (blue) respectively. The significant contributions from protein oxygen are marked with corresponding protein residue IDs containing the oxygen. The transparent background shows the pore profile within the open-state RyR1 structure, and the narrow pore region is highlighted between the black dashed lines. (b) Evolution of the z coordinates when Ca^2+^ ions (in different colors) permeate through the RyR1 channel from a typical segment of simulated trajectory. The occupied chamber accommodating either the selectivity filter or gate is shaded in gray unless both of them are occupied, which is indicated by green bars depicted with small arrows. Representative configurations with one of the chambers occupied by a Ca^2+^ ion (in silver) are shown above.

### The Ca^2+^ permeation pattern

The narrow region of the channel consists of the lower selectivity filter, the cavity, and the gate, which can be divided into two chambers by the saddle points of the Ca^2+^ density, as indicated by the dashed lines in Fig. 3a. The upper and lower chambers contain the binding sites S and G, respectively. The typical permeation pattern of Ca^2+^ ions is shown in Fig. 4b, from which it can be seen that permeation through the narrow pore region of the channel occur mainly in a one-at-a-time manner, meaning that there is only one Ca^2+^ residing in this narrow pore region, either in the upper or the lower chamber. The probability of both chambers being occupied by Ca^2+^ was only 2.4% in the trajectories, while the probability of only one chamber being occupied was 68.6%. Therefore, most of the time, only one Ca^2+^ can occupy the narrow pore region when permeating through the open-state channel. This permeation pattern is distinct from K^+^ and Na^+^ channels, in which usually both the narrow selectivity filter and cavity can be occupied by multiple permeating ions at the same time^11,13,16^. This is probably due to the fact that the electrostatic repulsion is much stronger between divalent ions than monovalent ions, and that RyR1 has a much shorter narrow selectivity filter region than typical K^+^ and Na^+^ channels (Fig. 4a).

## Discussion

The interaction between Ca^2+^ and protein is of great importance in studying Ca^2+^-mediated biological processes. Although the multi-site ion model cannot really represent the charge transfer and polarization effect explicitly, the simulation results showed that our new model behaves much better than conventional single-point ion models. Moreover, our model is entirely consistent with the currently widely used non-polarizable force fields, such as CHARMM and AMBER, and therefore can be easily used in MD simulations. As shown in the result section, our seven-site Ca^2+^ model can reproduce the solvation properties of Ca^2+^, including the hydration energy, the first solvation shell size (first-peak position of RDF), the coordination number, and the residence time of water in the first solvation shell, as well as the Ca^2+^-protein binding energies in a more quantitative way that is comparable to quantum chemistry calculations. The optimized Ca^2+^ model was validated by simulations of the RyR1 ion channel. In contrast to the conventional Ca^2+^ model, where ions get stuck in the channel in MD simulations, our new model resulted in the continuous permeation of Ca^2+^ through the channel, and the calculated conductance is in good agreement with the experimental electrophysiology result. Therefore, we believe this multi-site Ca^2+^ model can be widely used in simulating many Ca^2+^-involved bio-systems in addition to ion channels, and our simulations of RyR can provide detailed information about the Ca^2+^ permeation mechanism.

The ion binding and permeation mechanisms have been widely studied for K^+^ and Na^+^ channels with molecular dynamics simulations^10–12,14–17^, but only rarely studied for Ca^2+^ channels with conventional ion models^28^. From our MD simulations with the new multi-site Ca^2+^ model, the major Ca^2+^ binding sites in the open-state RyR1 were determined to be near the D4900 and G4894 residues on the luminal side, and the rate-determining step of permeation is found to be the step of passing through the selectivity filter constriction, where Ca^2+^ dwell for a relatively long time before passing through (Fig. 4b). Our simulation indicated that Ca^2+^ can easily accumulate in the wide upper selectivity filter, as shown by the high Ca^2+^ density from our equilibrium simulations (Fig. 3a). On average, there were about five Ca^2+^ or eight K^+^ in the selectivity filter in the presence of a 100-mv transmembrane potential (Fig. S1). This indicates that K^+^ cannot fill up the electrostatic energy well in the selectivity filter as effectively as Ca^2+^ do, which agrees with the previously proposed selectivity mechanism of the charge-space competition^32^.

The Ca^2+^ permeate through the narrow pore region of the channel following a one-at-a-time manner (Fig. 4b), which is distinct from the well-studied “knock-on” mechanism in K^+^ and Na^+^ channels. Previous studies have shown that multiple monovalent ions can enter the narrow pore region of the K^+^ and Na^+^ channels, and line up to facilitate the so-called ‘knock-on’ permeation, either tightly or loosely coupled^10–16^. It is not the case for the RyR channel, as only one Ca^2+^ was observed in the narrow pore region (r < 4 Å) during the Ca^2+^ permeation in our MD simulations, and this region contains both the lower selectivity filter and cavity. There may be two reasons for this different permeation pattern, one is that the narrow selectivity filter of RyR is shorter compared to that of K^+^ and Na^+^ channels (Fig. 4a), which can hardly accommodate multiple ions, and the second reason being that the electrostatic repulsion between divalent ions is much stronger than monovalent ions, which makes it energetically unfavorable for multiple divalent cations to sit side by side within a certain distance. In fact, we observed that when there was a Ca^2+^ in the cavity, other Ca^2+^ cannot enter the lower selectivity filter (Fig. 4b). Therefore, this unique one-at-a-time permeation pattern observed in our MD simulations occurs due to both structural features of RyR and strong repulsive interactions of divalent cations, which was not observed in previously studied ion channels.

Another distinct permeation feature of RyR is that Ca^2+^ ions are nearly fully hydrated during the translocation along the narrow pore region. Previous studies have shown that K^+^ is nearly fully dehydrated^11,16^ and Na^+^ is partially dehydrated^13,14^ during permeation, meaning all of the water molecules or at least several water molecules within the first solvation shell of the ions must be removed or replaced by other residues at the narrowest constriction sites. In fact, this dehydration process was believed to be the key factor responsible for the ion selectivity of the channels^13,17,33^, as different ions have different sized solvation shell and selectivity filters with different steric and chemical features can discriminate them by the free energy difference during the dehydration process. Interestingly, we did not observe such a dehydration behavior when Ca^2+^ was permeating through the RyR1 channel. As shown in Fig. 4a, the number of water molecules are nearly constant in the narrow pore region and the oxygen atoms coordinating with the Ca^2+^ from the protein is nearly zero throughout, indicating that the first solvation shell of the Ca^2+^ is intact during permeation, and no dehydration occurred when the ion passing through the narrow constriction sites in the selectivity filter and lower gate. This finding provides a clear picture on the exact hydration states of Ca^2+^ ions as they pass through the pore, which is consistent with earlier speculation that ion dehydration may not be a significant component of selectivity or permeation^34^. From the structural point of view, the open-state RyR1 is wider than K^+^ and Na^+^ channels in the selectivity filter region (Fig. 4a), and therefore sterically allows the Ca^2+^ to permeate with its first solvation shell intact. On the one hand, this allows highly efficient Ca^2+^ permeation to regulate ion concentration in the cytosol for muscle contraction and heartbeat, which is otherwise hard to imagine as Ca^2+^ has a much higher dehydration energy compared to K^+^ and Na^+^. On the other hand this probably also leads to the weaker ion selectivity of RyR1, compared to the highly selective KV, NaV, and CaV channels, since the powerful dehydration-based selectivity mechanism cannot be utilized here.

In summary, we developed a new multi-site Ca^2+^ model which behaves well when studying its interactions with proteins. It is entirely consistent with the widely-used CHARMM force field, and we are expanding to other force fields and divalent ions as well. With the new Ca^2+^ model, we revealed the detailed Ca^2+^ permeation process through the open-state RyR1, and discovered that multiple Ca^2+^ ions can accumulate in the wide upper selectivity filter, and then permeate through the narrow pore region following a one-at-a-time pattern, in which the permeating Ca^2+^ is fully hydrated with an intact first solvation shell (Fig. 5), distinct from the widely studied K^+^ and Na^+^ channels. These permeation details shed lights on the high efficiency and weak cation selectivity of the Ca^2+^ permeation through the RyR channels.

**Figure 5.**
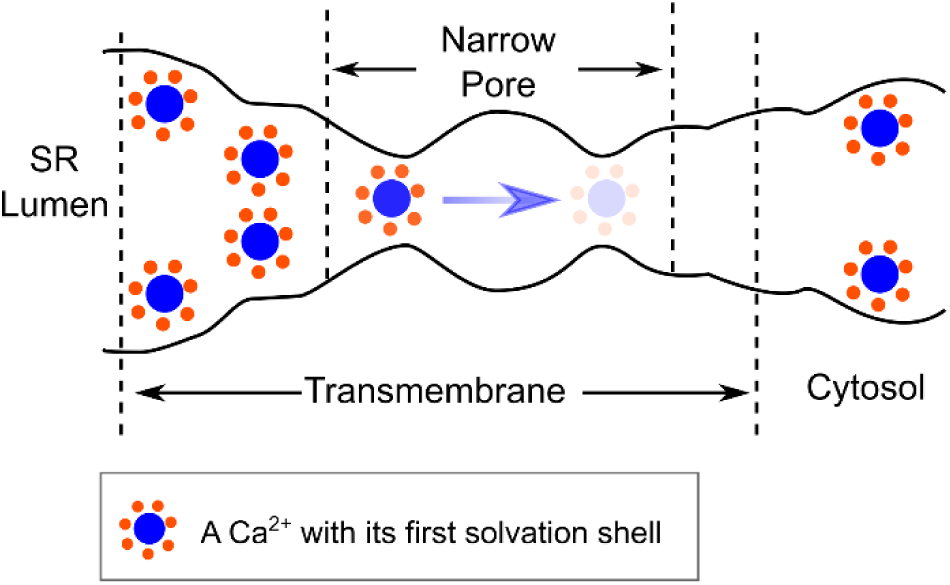
A sketch of the Ca^2+^ permeation mechanism of RyR1. There is only one Ca^2+^ in the narrow pore region, with a longer residence time at the selectivity filter constriction site (darker color), and a shorter residence time at the gate constriction (lighter color).

## Material and Methods

### I. Model Optimization

#### a. Simulations for Parameterization

All the MD simulations for the ion model parameterization were performed with OpenMM (version 7.0.1)^35^, as the package is highly flexible and can be easily customized with a Python interface, and the CHARMM force field^36^ (source: toppar_c36_aug15.tgz) was utilized in this work.

We built an ion-in-water system to optimize the ion properties in the water, which consists of one rigid multi-site Ca^2+^ ion in a cubic water box of 3 × 3 × 3 nm^3^. The TIPS3P water model was used in consistency with the CHARMM force field. In our simulations, NPT ensembles were generated by integrating the Langevin dynamics with a time-step of 2 fs and a collision frequency of 5 ps^-1^. The temperature was maintained at 298 K, and the pressure was regulated at 1 bar using a Monte Carlo barostat. Water molecules were kept rigid during simulations, and the cutoff of non-bonded interactions was 1 nm. For calculations of the radial distribution function, coordination number, and residence time, a 20-ns trajectory was generated. For hydration free energy calculations, 1-ns trajectories were generated for each of the 14 alchemical states (please also see below). The hydration free energy was estimated using the python implementation of the multistate Bennett acceptance ratio downloaded from https://github.com/choderalab/pymbar. The properties related to the radial distribution function were calculated using MDTRAJ^37^.

To optimize the ion-protein interactions, we used the Ca^2+^-Protein systems previously investigated by Li et al.^9^, and used OpenMM and CHARMM force field to perform the calculations as well (please see below for details).

#### b. Fitness Functions for Optimization

We define two fitness functions to optimize the parameters of our multi-site Ca^2+^ model. One is designated as the protein-fitness function 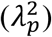 and the other as the water-fitness function 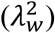. The protein-fitness function is defined against a dataset of quantum-mechanically calculated Ca^2+^-protein binding energies, 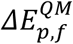, where *p* identifies the index among *N*_*p*_ (= 10) proteins, and *f* indexes *N*_*f*_ (= 21) trajectory snapshots for each protein. The formula for 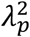 is

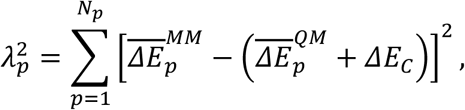

where 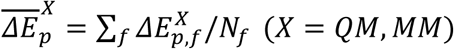, and *MM* indicates the molecule-mechanical results. Due to limitations of the methodology level and the basis-set size used in quantum-mechanical calculations, a systematic correction of binding energies (*ΔE*_*C*_ = 10 kcal/mol) is added to 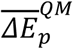 following Li et al.’s strategy^9^.

In addition to Ca^2+^-protein interactions, we also optimize our model to reproduce its energetical, structural, and dynamic properties in water. Specifically, these properties include the hydration free energy (*ΔG*_*h*_), the first-peak position of radial distribution function (*R*_1_), the coordination number (*N*_*C*_), and the residence time of water in the first coordination shell (*τ*_*R*_). The water-fitness function is defined as

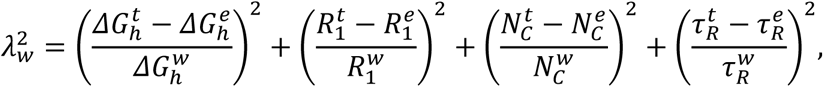

where quantities with superscripts of *t, e*, and *w* stand for theoretical, experimental, and weighting values, respectively.

#### c. Target Properties of Ca^2+^ in Water

We followed the approach of Mamatkulov et al.^38^ to determine the target hydration energy of Ca^2+^ (−1504 kJ/mol) as

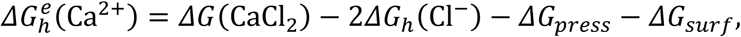

where *ΔG*(CaCl_2_) is the measured hydration energy of CaCl_2_ ^39^, *ΔG*_*h*_(Cl^−^) the theoretical hydration energy determined from Smith-Dang parameters^40^ for Cl^−^, *ΔG*_*press*_ the energy needed to compress one mole of ion gas at 1 atm into a liter, and *ΔG*_*surf*_ the energy change of crossing the air-water interface for 1 mol of ions. Marcus assessed the Ca-O internuclear distances in calcium salt solutions from different studies and concluded that the generally consistent result is 0.242 nm^41^, which was used as our target value for 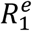. A recent neutron diffraction study^42^ revealed that the average number of water molecules in the first hydration shell of Ca^2+^ is close to 7, which we took as the target value for 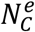. It is difficult to experimentally determine the residence time of water molecules in the first hydration shell of the calcium ion^43^, and a nuclear magnetic resonance (NMR) study estimated its value to be less than 100 ps^44^. Since the existing Ca^2+^ models generally overestimate *τ*_*R*_, we set 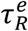 as zero in the course of optimization and checked the final result so as to be the same order of magnitude as 100 ps. The numerical values for the targeted experimental properties and their corresponding weights are listed in the second and third rows of Table S1, respectively.

#### d. Target Binding Energies of Ca^2+^ with Proteins

We followed Li et al.’s protocol to calculate the binding energies between Ca^2+^ and proteins^9^. As noted by Li et al., the default single-site model of Ca^2+^ in the CHARMM C36 force field generally overestimates the Ca^2+^-protein binding energies by ∼150 − 200 kcal/mol with respect to quantum-mechanical binding energies. They selected 10 high-resolution crystal structures of Ca^2+^-binding enzymatic proteins and performed MD simulations of the solvated proteins. From the equilibrated trajectories, 21 conformation snapshots were extracted for each protein and truncated to include atoms within a sphere of ∼0.55 nm around the ion. Quantum mechanical calculations were then carried out with the truncated models to obtain a dataset of binding energies, which is used in this work to optimize the multi-site Ca^2+^ model. For the systematic correction of binding energies (*ΔE*_*C*_) that is added to 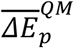, we use an estimation of 10 kcal/mol as Li et al. did in their work.

#### e. Calculation of Properties from Simulations

The theoretical hydration energy, 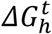, consists of two terms, i.e.

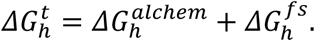

The first term refers to the free energy change corresponding to alchemically switching off the ion-water electrostatic and LJ interactions, and the second term is a correction due to the finite-size simulation box. It took ten and four steps to switch off the electrostatic and LJ interactions in our MD simulations, respectively. MD trajectories (1 ns) of one ion in a cubic water box (3 nm) were used to estimate the alchemical free energy by the method of multistate Bennett acceptance ratio^45^. For the finite-size correction, we took the formula derived by Hummer et al.^46^,

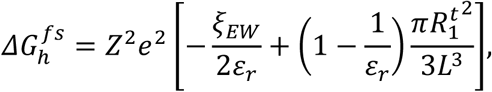

where *Ze* is the ion charge, *ξ*_*EW*_ is the Wigner potential, *ε*_*r*_ (= 82) is the relative dielectric constant, and *L* (= 3 nm) is the box size.

The first-peak position of the radial distribution function 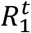 was calculated using the MDTRAJ software^37^ from a 20-ns trajectory, and the coordination number 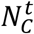 was computed by the integration of the first peak. We followed the definition described by Impey et al.^47^ to calculate the residence time 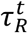. First, the residence time distribution *n*_*ion*_(*t*) was computed as

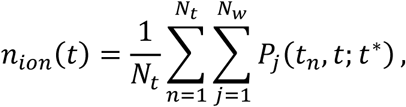

where *N*_*t*_ is the number of time frames and *N*_*w*_ is the number of water molecules. *P*_*j*_(*t*_*n*_, *t*; *t*\*) takes a value 1 if the water molecule *j* stays in the first hydration shell from *t*_*n*_ to *t*_*n*_ + *t* without leaving for any period larger than *t*\*, and takes the value 0 otherwise. Then, *n*_*ion*_(*t*) was fitted with 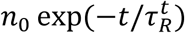 to obtain 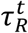.

The molecule-mechanical Ca^2+^-protein binding energies 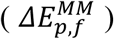 were simply calculated as the potential energy difference between Ca^2+^-bound and Ca^2+^-free configurations. Extra effort is required to first minimize the potential energies of Ca^2+^-binding configurations with respect to the ion’s orientation, since the multi-site model has lower symmetry than a single-point model.

#### f. Optimization Strategy

First, we did a thorough scan and prescreening, using the conventional optimization routines (such as conjugate gradient and basin hopping), random sampling, and sifting the parameter space by the properties described above in sequence. Initially, we focused on six parameters (*b*_*CD*_, *Q*_*C*_, *ε*_*C*_, *σ*_*C*_, *ε*_*D*_, *σ*_*D*_), i.e., treating interactions with water and protein on an equal footing. The searched parameter space as shown in Table S2 is determined from previous works^48,49^, and we substantially extended the range of *Q*_*C*_, *ε*_*C*_, and *ε*_*D*_ based on our optimization results. However, after systematic optimization, the calculated water properties and protein binding energies were still slightly off from the target values. Therefore, we further split (*ε*_*C*_, *σ*_*C*_) into 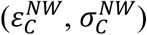 and 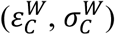, thus calculating the interactions of Ca^2+^ with protein and water separately. In fact, we took a two-step strategy to optimize our multi-site model. First, the water-fitness function 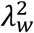 was minimized to obtain optimal values for 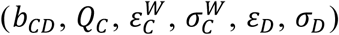. Second, with the parameters of (*b*_*CD*_, *Q*_*C*_, *ε*_*D*_, *σ*_*D*_) fixed, the protein-fitness function 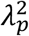 was scanned with a two-dimension grid to find the best values of 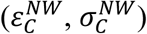 that reproduce quantum-mechanical Ca^2+^-protein binding energies. Due to the roughness of 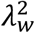 owing to the stochastic nature of the calculation, we developed a simple yet effective steep-descent-like algorithm that alternatingly searches along each dimension of the parameter space instead of the gradient. The parameter searching along each dimension was performed at linearly spaced steps, which were then finely tuned near the local minimum of a finite interval centering at the current minimal point. We used this strategy to further optimize the top parameters obtained in the initial stage of searching, which indeed yielded more satisfactory results. We found that the minimization mainly took place in the dimensions other than 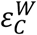 and *ε*_*D*_, which indicated that the landscape of the water-fitness function is much smoother along these two dimensions. Therefore, we selected a couple of combinations of 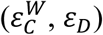 based on the scanning results and carried out further optimization until the fitness value was sufficiently low.

### II. Details of Permeation Simulation

To study the permeation of Ca^2+^ through RyR1, we performed MD simulations with GROMACS^50^ version 5.1.3 with the CHARMM36 force filed^36^ and the TIPS3P water model. The channel pore domain (residue 4820-4956) of the cryo-EM structure of the opened RyR1^21^ (PDB ID: 5TAL) was used as the starting structure. The *OPM*^51^ web server and Membrane Builder in CHARMM-GUI^52^ were used to build four POPC-RyR1 simulation systems.

A POPC-RyR1 simulation system with about 150 mM of calcium ions and a POPC-RyR1 simulation system with about 250 mM of potassium ions were built to calculate the conductance of the calcium ions and potassium ions with a transmembrane potential of 100 mV. Three 500-ns trajectories were conducted for the former system, and three 300-ns trajectories for the latter.

Another POPC-RyR1 simulation system with about 150 mM calcium ions was built to calculate the conductance of the calcium ions with our new calcium model with the same transmembrane potential of 100 mV. As the ion model cannot be simulated as a rigid body with Gromacs, we used bond and angle restraints to make the multi-site Ca^2+^ model as rigid as possible, which affect the thermodynamics of the Ca^2+^ to a minor extent. Six independent 500-ns trajectories were conducted. And, one 800-ns trajectory was run for the same system but without an electric field to obtain the stable binding sites of the calcium ions in the open-state RyR1.

All the simulation systems were first equilibrated with the standard CHARMM-GUI equilibration protocol followed by the production simulations with the position restraints applied on the *α* carbon of the protein (with a force constant of 1000 kJ·mol^-1^·nm^-2^). For all the production simulations, the periodic boundary conditions were used and the time step was 2 fs. The v-rescale algorithm with a time constant of 0.5 ps was used to maintain the temperature at 310 K, and the Parrinello-Rahman algorithm^53^ with a time constant of 1 ps was used to maintain the pressure at 1.0 bar. The Particle-mesh Ewald method^54^ was used to calculate electrostatics, and the cut-off length of the van der Waals interaction was 1.2 nm.

## Acknowledgement

We thank Prof. Sergei Noskov and Prof. Roland R. Netz for data sharing and discussions with us. The research was supported by grants from the Ministry of Science and Technology of China (National Key Research & Development Program of China, 2016YFA0500401), the National Natural Science Foundation of China (grant no. 21873006), and the Young Thousand Talents Program of China. Part of the molecular dynamics simulation was performed on the Computing Platform of the Center for Life Sciences at Peking University.

## Supplementary Information

**Table S1.**
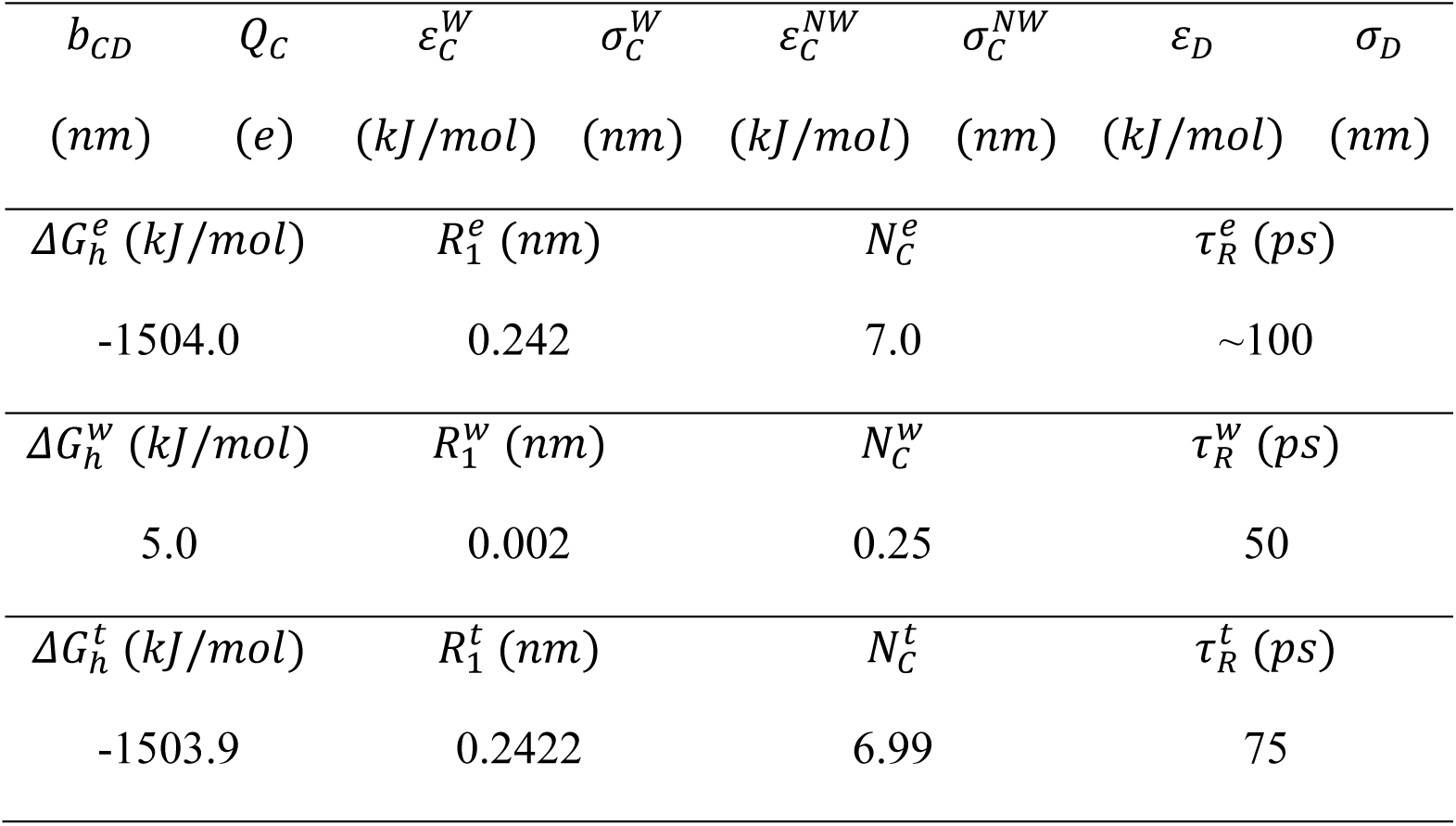
The optimized parameters (1^st^ row), target experimental properties (2^nd^ row), property weights (3^rd^ row), and theoretical values (4^th^ row) calculated using the optimized parameters.

| $b_{CD}$ | $Q_C$ | $\varepsilon_C^W$ | $\sigma_C^W$ | $\varepsilon_C^{NW}$ | $\sigma_C^{NW}$ | $\varepsilon_D$ | $\sigma_D$ |
| --- | --- | --- | --- | --- | --- | --- | --- |
| (nm) | (e) | (kJ/mol) | (nm) | (kJ/mol) | (nm) | (kJ/mol) | (nm) |
| $\Delta G_h^e$ (kJ/mol) | | $R_1^e$ (nm) | | $N_C^e$ | | $\tau_R^e$ (ps) | |
| -1504.0 |  | 0.242 |  | 7.0 |  | ~100 |  |
| $\Delta G_h^w$ (kJ/mol) | | $R_1^w$ (nm) | | $N_C^w$ | | $\tau_R^w$ (ps) | |
| 5.0 |  | 0.002 |  | 0.25 |  | 50 |  |
| $\Delta G_h^t$ (kJ/mol) | | $R_1^t$ (nm) | | $N_C^t$ | | $\tau_R^t$ (ps) | |
| -1503.9 |  | 0.2422 |  | 6.99 |  | 75 |  |

**Table S2.** The parameter space scanned for the optimization of the multi-site Ca^2+^ model.

| $b_{CD}$ | $Q_C$ | $\varepsilon_C$ | $\sigma_C$ | $\varepsilon_D$ | $\sigma_D$ |
| --- | --- | --- | --- | --- | --- |
| (nm) | (e) | (kJ/mol) | (nm) | (kJ/mol) | (nm) |
| [0.05, 0.15] | [-8, 2] | [0.1, 10.1] | [0.20, 0.32] | [0.1, 10.1] | < 0.01 |

**Figure S1.**
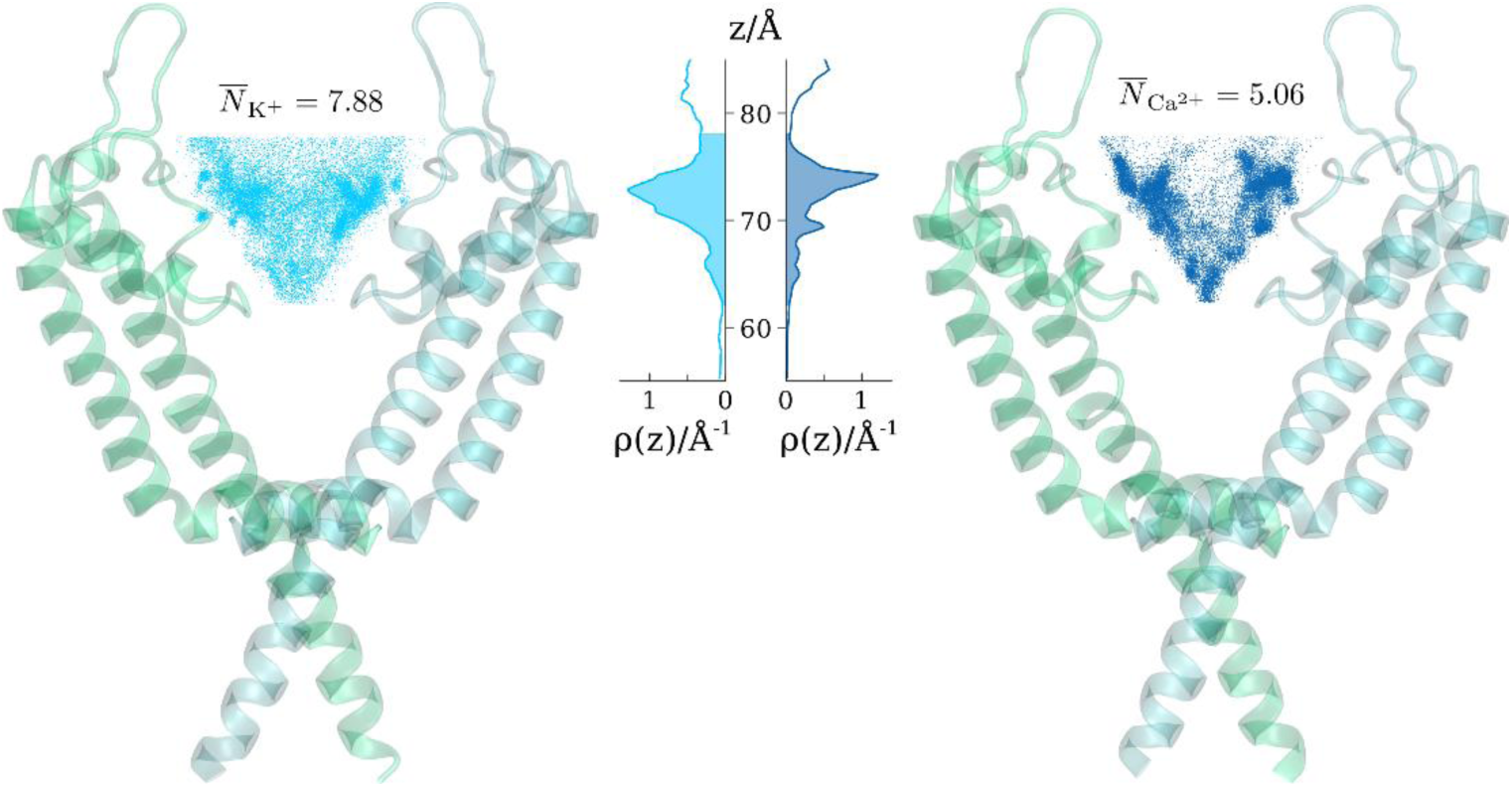
The average number of ions occupying the selectivity filter of RyR1. The number was calculated for K^+^ (left) or Ca^2+^ (right) in the presence of a 100-mv trans-membrane potential. The ions located above the selectivity filter constriction site and within the pore were considered. The positions of the ions, represented by dots in the protein structures, were sampled from 6 × 500-ns trajectories (Ca^2+^) and 3 × 300-ns trajectories (K^+^) with a ratio corresponding to their average occupying number. The ion number densities along the channel axis were given in the middle panel, and the integration of the shaded area yielded the average number of ions within the filter. In this figure, K^+^ and Ca^2+^ are referred to by the cyan and blue colors, respectively.

